# The dynamics of phage predation on a microcolony

**DOI:** 10.1101/2023.06.13.544539

**Authors:** Rasmus Skytte Eriksen, Frej Larsen, Sine Lo Svenningsen, Kim Sneppen, Namiko Mitarai

## Abstract

Phage predation is an important factor for controlling the bacterial biomass. At face value, dense microbial habitats are expected to be vulnerable to phage epidemics due to the abundance of fresh hosts immediately next to any infected bacteria. Despite this, the bacterial microcolony is a common habitat for bacteria in nature. Here we experimentally quantify the fate of microcolonies of *Escherichia coli* exposed to virulent phage *T4*. It has been proposed that the outer bacterial layers of the colony will shield the inner layers from the phage invasion and thereby constrain the phage to the colony’s surface. We develop a dynamical model that incorporates this shielding mechanism and fits the results with experimental measurements to extract important phage-bacteria interaction parameters. The analysis suggests that, while the shielding mechanism delays phage attack,*T4* phage are able to diffuse so deep into the dense bacterial environment that colony level survival of the bacterial community is challenged.

**Significance:** The shielding mechanism of bacterial microcolonies against phage invasion has been experimentally quantified, and the result was analyzed by using a mathematical model. The analysis suggests that while forming a dense colony delays phage attack by limiting the infection closer to the colony surface,*T4* phage can diffuse so deep into the dense bacterial environment that colony-level survival of the bacterial community is challenged. This finding highlights the importance of the interplay between the phage infection dynamics and the physical structure of bacterial colonies in controlling the microbial population.

## Introduction

Bacteria often live in spatially structured environments e.g. in soil[1, 2], in solid foods[3], in the human gut microbiome[4, 5], and in biofilm[6, 7, 8]. Even in liquid conditions, bacteria are often arranged into small clusters[9]. These structured environments change the interaction between the bacteria and their phage predator compared to the well-mixed conditions [6, 10, 11, 12, 13, 14] which are typically covered by mathematical models [15, 20, 21, 22, 23, 24, 25, 26, 27, 16, 17, 18, 19].

We have proposed that a microcolony exposed to phages can, in some cases, grow faster than the phages can kill the bacteria in the colony[11]. This is due to an intricate emergent structure formed by the invading phages, which causes cell lysis to occur primarily on the surface of the bacterial colony. The mechanism that causes this structure can be understood by focusing on a single phage infection cycle: The phage invasion begins with external phage particles reaching the microcolony, causing bacteria on the surface of the colony to be infected by phages. While the phage infection in the surface bacteria matures, more external phages will reach the colony. The high phage adsorption rate, and the fact that infected bacteria continue to adsorb phage particles, limits the phages from penetrating more than a few layers into the colony. This means that most phages will hit bacteria that are already infected and only few phages will penetrate sufficiently deep to hit an uninfected bacterium and cause a new infection cycle. Due to the decreasing chances of these infections, they occur later than the infections nearer the surface. This causes a gradient in the timing of the infections, where bacteria nearer the surface is closer to lysis. Once the surface infections have matured to lysis, most of the released phages will hit the nearby bacteria which are already infected, and, like the original phage arriving to the colony, only few phages will penetrate sufficiently deep to reach uninfected bacteria.

This clear separation of the infected bacteria on the surface of the colony and the uninfected bacteria nearer the core suggests that if the colony is large enough, then the growth of the core bacteria, which is proportional to the radius cubed, can be larger than the killing by phage, which is proportional to the radius squared.

In a previous experiment[11], we showed evidence that suggested the existence of such a core of uninfected bacteria. This was further substantiated by a clear transition from small colonies being quickly eliminated by the phages to larger colonies growing to substantial size despite the killing by phage.

What is not fully clear from this experiment is whether the bacterial colony is indeed able to continuously grow despite the phage attack, or if we instead observe the remains of a slowly declining bacterial population. Furthermore, the previous experiment considered a virulent mutant of the temperate phage *P1* (*P*1 _*vir*_) rather than a naturally strictly lytic phage. This difference may be important since lytic phages typically have substantially shorter latency times and larger adsorption rates[28]. Our previous results suggest these parameter differences may work against each other: The shorter latency time should increase the virulent potency of the phage, while the increased adsorption rate should, somewhat counter-intuitively, reduce the killing by phage by preventing the phage from reaching far into the microcolony.

In this work, we use the lytic phage *T*4, which has a latency time of *τ*_*L*_ ∼ 23 min. This is roughly one third as long as for *P*1 and phage *T*4 has roughly twice the adsorption rate of *P*1[28]. In terms of dynamics, the change in latency time is expected to have a larger impact than the changes to the adsorption rate. This is highlighted by the model prediction of the “critical radius”, *R*_*c*_, in Ref. [11]:

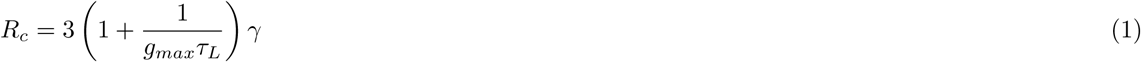

where *g*_*max*_ is the maximal bacterial growth rate and *γ* is the phage penetration depth (See Fig. 1 for definition of *γ*). The critical radius indicates the minimal size a colony needs to be in order to outgrow the killing by phage. Notice that this expression decreases with the latency time. Conversely, the adsorption rate of the phage influences *γ* in a non-linear way, which only slowly changes at high adsorption rates [11]. The *T*4 phage is also different from *P*1 phage in another way, namely, the phage is known to cause “lysis inhibition”[29, 30] which is a process whereby the number of co-infecting phages, the so-called “multiplicity of infection” (MOI), can change the lytic cycle into a second mode of infection where the latency time is substantially longer but with a correspondingly higher phage burst size.

**Figure 1.**
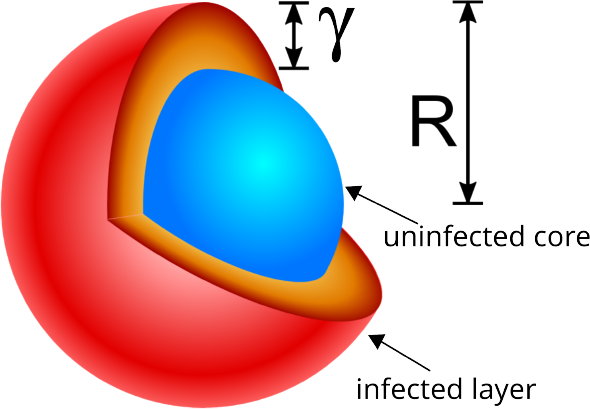
Illustration of model principle. Our model divides the colony into two parts: an outer shell (shown in red colours) of thickness *γ*, which is being preyed upon by the phage population, and an inner core (shown in blue colours), which is phage-free and contributes to the growth of the colony. The total colony radius *R*and the phage penetration depth *γ* describe the model colony.

In this paper, we expand on our previous work in a number of ways to challenge the proposed model of a core and shell structure for phages invading bacterial microcolonies. In terms of the experimental design, we use a phage with markedly different properties and behaviour and couple this with more detailed measurements of the dynamics of phage invasion. This last part is achieved by the use of a microscope with a motorized XY scanning stage that allowed us to track the time evolution of several individual colonies of different size as they are preyed upon by phages. We combine these experimental measurements with a dynamical model of a microcolony based on the core-shell structure phage invasion to analyze if the model can reproduce the behaviour and exact essential parameters such as *γ*. We find that the phage T4 is unexpectedly efficient in penetrating deep into the colony and killing the bacteria.

## Materials and Methods

### Experiments

#### Infection of *Escherichia coli* microcolonies by T4 bacteriophage

*E. coli* strain SP427[31], an MC4100 derivative containing chromosomally encoded *P*_*A1/O4/O3*_*::gfp*mut3b gene cassette[32] was grown overnight in LB and then diluted to 10^3^ CFU*/*mL in MC buffer(50 mM CaCl_2_ and 25 mM MgCl_2_). 10μL of the diluted overnight culture were mixed with 1 mL of soft agar (10 g tryptone, 8 g NaCl, 5 g yeast extract, and 5 g agar for 1 L. 0.2 % glucose, 50 mM CaCl_2_, 25 mM MgCl_2_, 10 mM Tris-HCl pH 7.4 were added after autoclaving) and poured into a well in a 6-well glass-bottom plate (Cellvis P06-1.5H-N, well diameter 35 mm). This yielded a soft agar layer with a thickness of about 1 mm. This process was repeated at 30-minute intervals for each well to provide an ensemble of colonies at various stages of development. In between the poring, the plate was kept in a 37 *°*C incubator. In the last round of plating, two wells were plated simultaneously with one serving as a control well. The plate was then incubated at 37 *°*C for 5 hours, after which the 5 non-control wells were infected with phage T4 by spraying 500μL of high titer (10^10^ PFU*/*mL) bacteriophage lysate onto each well using a perfume spray bottle. The plate was then placed under a microscope at 37 *°*C and brightfield and fluorescence images of the growing colonies were captured every 30 minutes.

### Image analysis

Our experimental time-lapse setup outputs a series of images in a bright-field and green fluorescence configuration. We have developed an algorithm to analyse these images that filters the image data and estimates the size of the bacterial colony. The algorithm works by exploiting the differences in luminance between the bacterial colony and the growth medium. The area *A* of the 2D colony cross section can be extracted from the image by applying a threshold value to the bright-field and green fluorescence images. Next, the algorithm assumes the colony is fully spherical and determines the corresponding colony radius 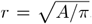. For the bright-field images, the algorithm is insensitive to the chosen threshold value since the colony outline has a large contrast with the soft agar growth medium. This is not so for the green-fluorescence signal, which shows a gradient from the centre towards the edges of the colony. When the infected bacteria lyse, they release their fluorescent proteins, which diffuse away over time but, on shorter time scales, still contribute to the green fluorescence signal. This means that the chosen threshold values effectively constitute an additional model parameter that must be treated with extra care (see Section S2 in the supplement). The analysis scripts are available as per the data availability statement.

### Mathematical model

We base our model on two previously published results. In Ref. [11], the authors show that phages invading a bacterial microcolony have a limited ability to penetrate into the colony. This is due to an emergent phenomenon whereby the time delay between infection and lysis causes the infected bacteria to be spatially structured in a gradient, where bacteria who are close to lysis are located towards the very outside of the colony and newly infected bacteria are located deeper in the surface layers of the colony. The phage progeny released by lysis at the colony surface thus has to move through layers of already infected bacteria to reach the uninfected bacteria nearer the colony’s centre. This prevents the phages from easily reaching the core of the colony. Meanwhile, the uninfected bacteria in the core divide to create new bacteria and may possibly overcome the loss mediated by phage predation.

Following the above observations, we model the colony as consisting of two parts. A growing core of uninfected bacteria is surrounded by a shell of thickness *γ* being attacked by phages (see Figure 1).

The growth of such a spherical bacterial microcolony has previously been studied in Ref. [33] in the absence of phages. Here, the authors realistically model how a three-dimensional bacterial colony grows when confined in a soft medium. Under the modelled conditions, the bacteria divide symmetrically in all directions and thus form a spherical colony. In order for the bacteria to grow, they consume the nutrients present in the soft agar medium. The bacteria consume the nutrients locally, and over time, this results in a nutrient density gradient, where all nutrients are consumed at the centre while fresh nutrients reach the outer layers of the colony by diffusion. This causes the colony to initially grow exponentially until the local nutrients are consumed, after which the colony grows in size at a roughly constant speed set by the diffusion rate of the nutrients.

We model the growth of the uninfected bacterial core of our bacterial colony in a similar way, but with some adjustments to allow for the modelling of the infected shell of bacteria. Our full model is described below, but essentially, we simplify the modelling of the nutrients such that we no longer explicitly model the density of the growth-limiting substrate but rather measure nutrients in units of cell divisions (one unit of nutrients yields one new bacterium). As a more technical change, we re-cast their equations onto an integro-differential form, which allows us to model the colony radius directly (this possibility was mentioned in the original paper).

Our analysis focuses on modelling the interactions between phages and a growing spherical bacterial colony. For such a spatially symmetric problem, it is ideal to use spherical coordinates since every variable will only change along a single coordinate *r*. We assume the cells are in the exponential growth phase, where the core of uninfected cells is assumed to grow at a rate of *g*(*n*). This growth rate is a function of the (local) density of available nutrients *n*, which controls the growth rate in a Monod-like way through the Michaelis constant *K* [33]:

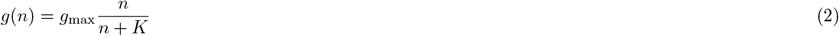

Notice that while not explicitly written in the notation, the nutrients are functions of space and may vary as a function of *r*.

The latency time between initial phage adsorption and the lysis of the host is important for the emergence of the shell-core structure of the bacterial colony. The relationship between this latency time and the metabolic state of the host bacterium varies between phage strains with some phages halting their reproduction as the nutrients are depleted [34] and others, e.g. *T*7, which can propagate even on stationary-phase bacteria [35]. For phage T4, the phage growth parameters such as burst size and latency time have been quantified in the steady state exponential growth in different growth media to quantify the host growth rate dependence of these parameters [34]. However, how the phage latency time and the burst size decrease as the bacteria enter the stationary phase is still unclear [36].

For our purposes, we use the growth rate to control the rate at which the attacking phages lyse their hosts (thereby assuming the phage reproduction rate scales with the depleting nutrients). The concentration of nutrients is our only real measure of how close the bacteria are to the stationary phase and, therefore, serves as a reasonable proxy for phage productivity. This means that in our model, the killing rate by phages is assumed to be proportional to the growth rate by a factor of *ϵ*.

Our simulated colony thus consists of two parts: the inner core of uninfected bacteria, which grow and divide, and the outer shell of bacteria, where the phages kill bacteria and penetrate the colony.

Notice that the nutrient level across the colony is expected to vary and change over time. To write an equation for the colony radius *R*, we must then consider the contributions of both parts of the colony and integrate over nutrients locally available to each part:

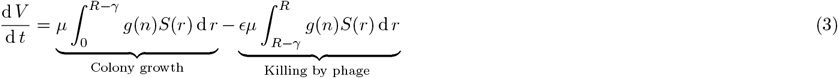

Where *S*(*r*) = 4*πr*^2^ is the surface area at a distance *r* from the center, 1*/μ* is the bacterial density in the colony, and *R* is the (total) radius of the colony. Since the right hand side of Equation (3) is dependent on the radius *R*, we rewrite the equation to give the time derivative of the radius instead of the volume.

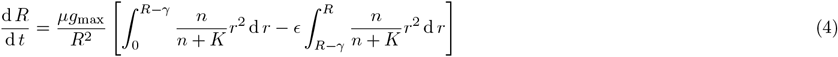

The nutrients in the system consist of small particles which we assume are unconfined in the medium. As such the nutrients can diffuse with a constant of diffusion constant *D*_*n*_, while the growing bacteria inside the colony are consuming the nutrients:

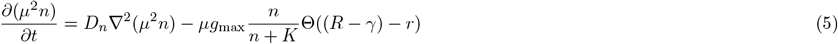

where ∇^2^ is the Laplacian and Θ(*x*) is the Heaviside step-function. This function is 0 if *x <*0 and 1 if *x >*= 0, which ensures nutrients are only being consumed inside the colony’s core (which has radius *R* − *γ*). The factor *μ*^2^ (a constant) is multiplied to the whole equation so that *g*_*max*_ here appears multiplied by *μ*.

In order to compare with experimental results, we model how the total biomass, dead or alive, increases in volume over time.

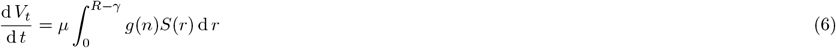

We deviate from the work in Ref. [33], by the use of the integro-differential equation instead of resolving the bacteria on a lattice. We prefer this solution for a number of reasons. First, it elegantly solves the issue of “cell shoving”, whereby bacteria have to be redistributed when solved on a lattice. This issue arises since the equations do not take the physical size of the lattice points into account but rather let the densities increase unrestricted which eventually leads to physically impossible densities of bacteria. Secondly, the integro-differential equations allow us to model *R* directly and since this variable enters all the equations it is much simpler to implement this way. Note that we still solve the nutrient density on a spatial grid since it is paramount that it varies across the colony.

We are interested in situations where the radius of the colony varies from a few μm initially to around 500μm toward the end of the simulation, In such situations, the simulation will need good spatial resolution around *r*= 0 μm where the colony is small, but as the colony grows the need for spatial fidelity decreases. Therefore, we choose to implement the spatial grid as a non-uniform grid where the first node is located at *r*_0_ = 0 μm, the second is located at *r*_1_ =Δ*r*, the third is located at *r*_2_ =*s*Δ*r* and the *k*’th is located at *r*_*k*_ =*s*^*k−*1^Δ*r*. Since this is a system without the influx of new nutrient, we impose the reflecting (von Neumann) boundary conditions on the nodes located at *r*_0_ and *r*_*N*_. The Laplacian is more difficult to implement on such a non-uniform grid so we have included a detailed derivation in the supplement (see S1).

The strength of the phage attack is determined by the killing parameter *ϵ* and the penetration depth *γ*. It is possible that the latter may change over time as the initial phage invasion takes hold and as the colony grows and thereby presents a larger target for the free phages.

There are several choices for how to model the penetration depth *γ*. Here we follow the model proposed in Ref. [11], by using a simple step function on the form:

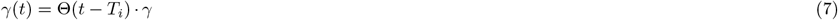

This model mimics the behavior where: 1) no phages are present before *t*=*T*_*i*_ and 2) the phage invasion penetrates a fixed distance *γ* into the colony.

As discussed in the Image analysis section, the threshold value chosen when analyzing the region of the colony expressing green fluorescence effectively constitutes an additional model parameter. Rather than implementing this threshold into the main modelling, we instead temporarily introduce the new model parameter *v*. We use this parameter to describe how the density of dead bacterial matter relates to the density of alive matter. In our model, the colony’s bright-field radius 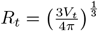 is derived from the volume has contributions from the debris from dead bacteria surrounding the central colony and from the alive colony of radius *R*. We now include our temporary density parameter *v* and redefine the bright-field radius as:

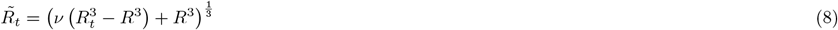

The experimental measurement of the fluorescent region varies substantially based on the chosen threshold value. We, therefore, fit our model to data corresponding to a range of threshold values. These fits, in turn, predict a range of values for *v* ranging from ∼ 0.2 to ∼ 1.4. Here we argue that the value *v* is expected to be around 1, meaning that dead bacteria, on average, take up the same space as alive bacteria. By choosing the threshold value corresponding to *v* ≈ 1 we obtain a self-consistent measure for the threshold value (see S2 in the supplement). We have also analyzed how the result changes with the threshold value for the alive region fluorescent signal (hence by changing the value of *v*), and the outcome is presented in the result section.

### Parameter fitting

We use the data from 5 colonies grown without phage application for estimating the growth parameters. We used the ensemble average *R*_*BF*_ (*t*_*i*_) and standard deviation *σ*_*BF*_(*t*_*i*_) of the alive radius calculated from the BF images at each observation time point *t*_*i*_. The growth parameters are estimated by minimizing the difference

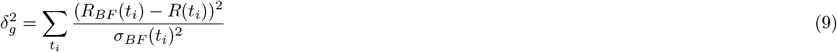

where the alive radius *R*(*t* _*i*_) is calculated from eq. (4) with setting phage attack parameters zero.

We fitted the nutrient diffusion constant *D*_*n*_, the initial nutrients *μ*^2^*n*, the maximum growth rate *μg*_*max*_, and the Monod constant *μ*^2^*K* (the latter three parameters appear with *μ* because we only need these combined parameters in the model). For *D*_*n*_, it has been reported that the small molecule diffusion constant is about 10^6^ μm^2^*/*h [37], but since nutrients consist of various molecules and we do not know what is rate-limiting for growth, we let it be one of the fitting parameters.

For fitting to the phage attack parameters (*ϵ* and *γ*), the alive radius *R*(*t*_*i*_;*T*_*j*_) calculated from eq. (4) and the total radius 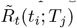 calculated from eq. (8) at measurement time *t*_*i*_ for a given phage attack timing *T*_*j*_ are used to calculate the distance between the data and the model fit

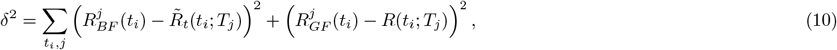

where the index *j* denotes the different colonies. Note that *T*_*i*_ is a fitting parameter that varies for each colony. Then, the phage attack parameters are estimated by minimizing the *δ* ^2^.

When calculating *R* and 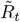, we used a non-uniform spatial discretization described in the model subsection, and the Matlab function *ode15s* was used to integrate the equation in time direction. The objective functions’ minimisations were done using Matlab function *fminsearch*. An early stopping criterion was added to minimise the phage attack parameters. The minimization was performed iteratively, where each iteration performed a number of minimization steps equal to 10 times the number of free parameters. When a minimization step improved the objective function by less than 5 percent, the minimization iteration stopped.

## Data availability statement

Our analyses and models are implemented in MATLAB version 2022b (9.13.0.2105380). We have made the full code and data available online[38].

## Results

### Imaging of growing and phage attacked colonies

In the experiment, we plated *E. coli* strain SP427 into all the wells of a 6-well plate with a glass bottom. Initially, a small number of cells are scattered and embedded in soft agar with nutrients (See Materials and Methods for detail). The soft agar used is dense enough to prevent long-distance swimming of bacteria (e.g. [39]). Therefore, cells grow locally to form spherical colonies. One of these wells was used as a phage-free control and the other five were eventually exposed to phages. In turn, one of the five wells were plated every half hour to create a time-staggered series for the five wells, with the control well being plated simultaneously with the last of the 5 wells. Between platings, the 6-well plate was incubated at 37 *°*C inside our enclosed, temperature regulated, inverted microscope. Once all of the wells were plated with bacteria, we identify the location of a total of 30 colonies divided equally between the six wells.

With the colony locations noted, we let the colonies incubate until 5 h has passed from the last plating. At this time, all but the control well is sprayed with the virulent *T*4 phage. Due to the time-staggered plating of the wells, the colonies have had between *T*= 5 h and *T*= 7 h to grow before the addition of phages.

The 6-well plate is then returned to the microscope for further incubation. The *E. coli* strain SP427 constitutively express the green fluorescent protein. We programmed the microscope to image the brightfield and green fluorescence expression of each of the 30 bacterial colonies every half hour over a period of 16 hours (see Figure 2 for an example). We then analyze the microscope images to determine the radius of the visible bacterial colony under bright-field conditions and the radius of the region visible under green fluorescence conditions.

**Figure 2.**
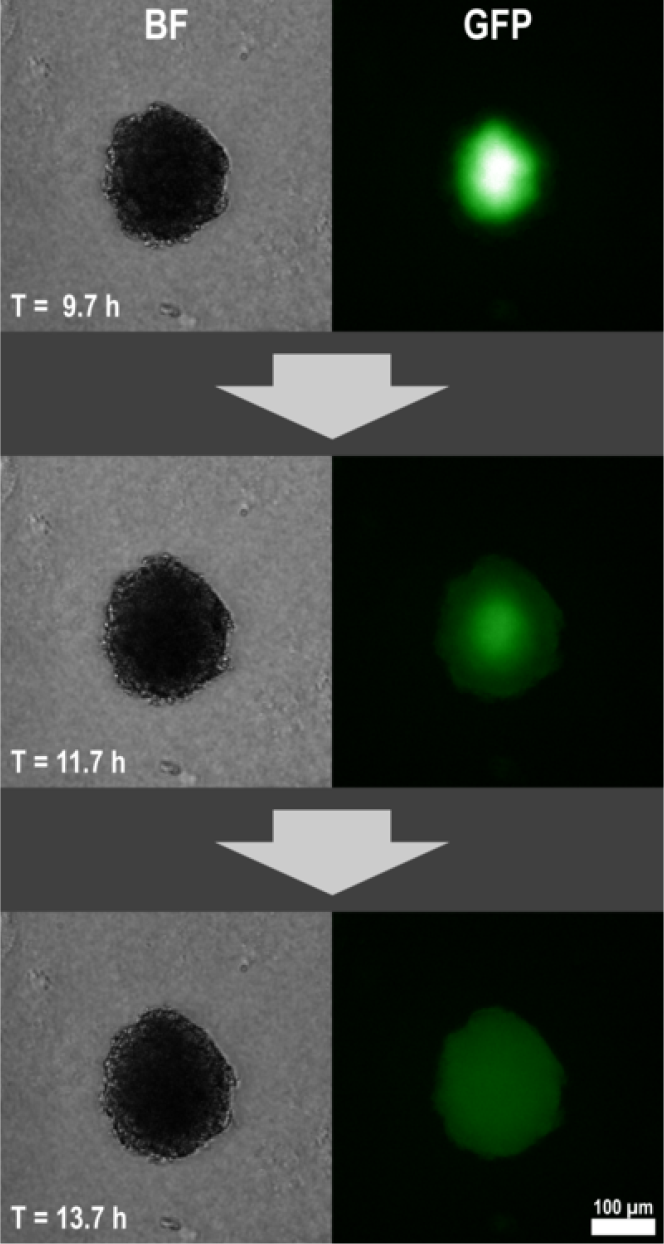
Structured killing by phage. We show snapshots of an *Escherichia coli* colony under attack by phage *T*4. The bright-field (BF) images show the total bacterial biomass while the signal from the green fluorescence protein (GFP) indicates the presence of alive bacteria. The phage invasion moves from the outside of the bacterial colony towards the centre until all bacteria in the colony are killed.

In Figure 2, we show images of a colony as the phages invade the colony. Under the brightfield (BF) conditions, the colony appears to change little indicating a distinct lack of growth. At the same time, the green fluorescence images show the loss of fluorescence starting from the colony’s surface, indicating the gradual elimination of the alive bacteria within the colony. We, therefore, used the fluorescent region as a proxy for the region of alive bacteria.

After image analysis, we obtained growth curves for both the total biomass and the alive biomass in the colony for all 30 colonies (See Materials and Methods for details). Of the 25 colonies which were sprayed with phage, 4 colonies exhibit either anomalous growth behaviour or growth behaviour consistent with the emergence of phage-resistant bacteria that keep growing even they are exposed to phages (see example in Figure 3). Our model does not include the resistant cells, but we discuss their effect later in the discussion. For one additional colony, we had difficulties detecting the green fluorescence region during image analysis. The growth curves of these colonies are included in the supplement (see Figure S3) but otherwise excluded from further analysis. In total, this leaves us with the trajectories of 20 phage-attacked colonies.

**Figure 3.**
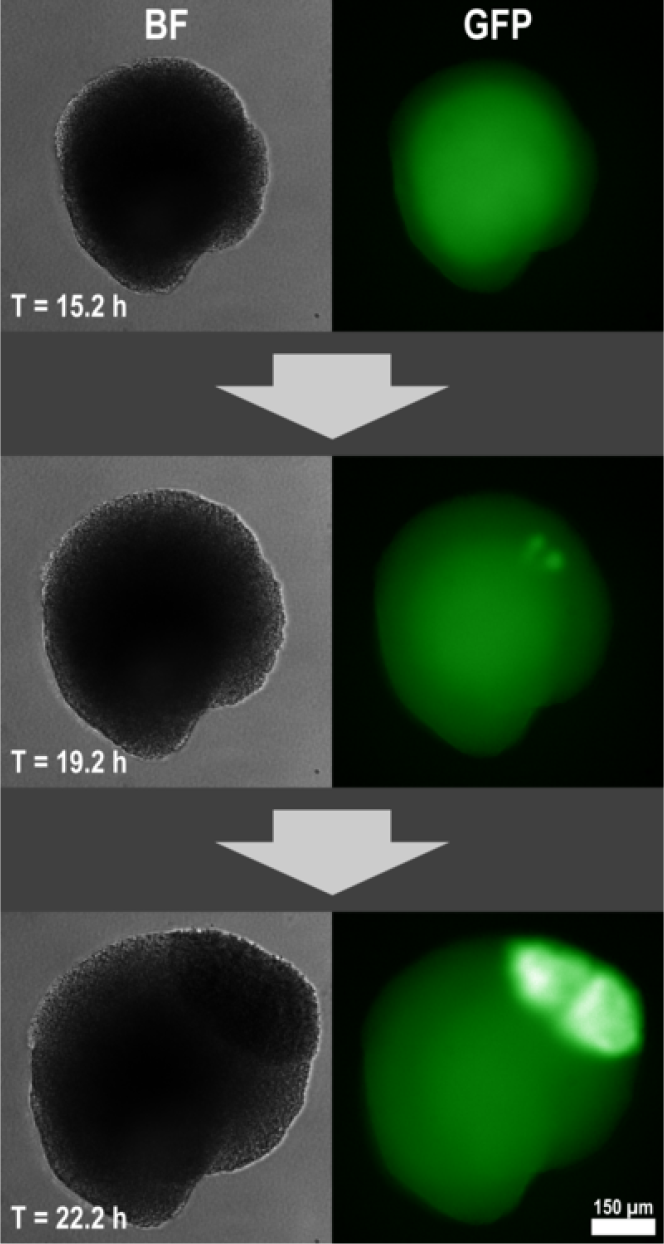
Emergence of resistant bacteria. These snapshots show a microcolony that is on the verge of total collapse as indicated by the faint GFP signal in the upper frame. However, small pockets of resistant mutants emerge and form small clusters of growing bacteria with a strong GFP signature.

### Growth curve fitting to the colonies without phage attack

From our time-lapse data, we obtained the growth curves for five control colonies not exposed to phages. In order to reduce the likelihood of overfitting our model, we use this data alone to fit the growth parameters of the model.

The model equations used for this fit are (4) and (5), with the phage penetration depth *γ* and the rate of killing by phage *ϵ* set to zero. In practice, it is difficult to disentangle the various factors controlling growth from a single batch growth curve[40]. We, therefore, need to restrict the model-fitting: In our framework, the two parameters *μ* (the volume occupied by one cell in a colony) and *g*_*max*_ (the maximum growth rate in the Monod-growth function eq. 2) always occur together in the form *μg*_*max*_, so we treat them as one parameter.

The model fit also contains the initial radius of the colony as a parameter. We set the initial volume of the colony, which is expected to start from a single cell, to 1.4 μm^3^ on average (hence a radius of 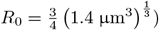), assuming that the newly born cell has a volume of 1 μm^3^ and a cell can be anywhere in the cell cycle with equal probability. In addition, we restricted the fitting to the first 16 h of growth. After this point in time, the bacterial colony has reached a size comparable to the thickness of the agar, which means the colony is likely to be no longer spherical and the nutrient field is not expected to be spherically symmetric anymore as assumed in the model.

The parameters are optimized by minimizing the difference between the experimentally obtained alive colony radius and the model-predicted alive radius (See Materials and Methods). The fitted growth curve with the estimated growth parameters is shown in Figure 4. It shows that the model captures the growth dynamics of the colony. Note that if we fit it against the entire dataset, the model will be able to describe the observed growth throughout the experiment.

**Figure 4.**
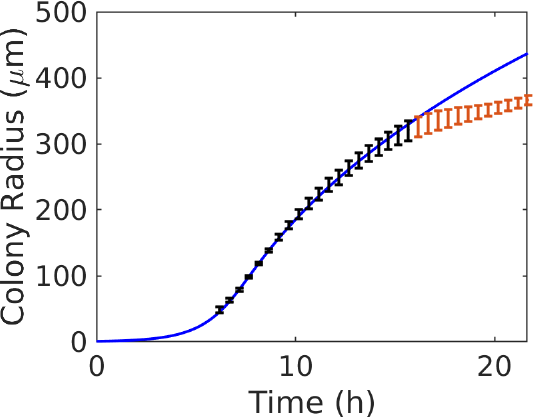
Growth curve of the bacteria. The 5 control colonies of *Escherichia coli* show undisturbed growth in our experimental setup. Black error bars indicate the standard error of measured growth. Red error bars indicate growth post 16 h that is excluded from the fit. The blue line indicates the least-square fit of our model eq. (4). See text for detail of the fit. The reduced *χ*^2^ value for the obtained fit was 0.21.

The maximum-likelihood fits provided us with the parameter values for our model along with the estimated uncertainty on these parameters. Due to the above-mentioned propensity for over-fitting of the growth parameters, we stop the fitting of the model once the fitting landscape is locally flat. Specifically, we stop fitting once any individual perturbation of parameters does not improve the fit. This also means that the parameter estimates should be considered cautiously. The full list of model parameters and their values is listed in Table S1 in the supplement. In Table 1 we provide an overview of the relevant growth parameters.

**Table 1:** Fitted growth curve parameters. We show the fit parameters and their standard deviation corresponding to the growth curve in Figure 4. These are presented in an easy-to-read format. For a full list of parameters see Table S1. Note that the volume of one bacterium in a colony, *μ*, appears in combination with other parameters and cannot be fitted independently. Hence, it is included as a part of fitted values in the above table and also affects their unit.

| Description | Value |
| --- | --- |
| Nutrient diffusion constant $D_n$ | $(1.29 \pm 0.03) \cdot 10^5 \mu\text{m}^2/\text{h}$ |
| Initial nutrients $\mu^2 n$ | $(4.68 \pm 0.09) \cdot 10^{-2} \mu\text{m}^3$ |
| Maximum growth rate $\mu g_{max}$ | $(3.33 \pm 0.012) \mu\text{m}^3\text{h}^{-1}$ |
| Monod constant $\mu^2 K$ | $(2.75 \pm 0.06) \cdot 10^{-2} \mu\text{m}^3$ |

### Fitting of the phage-related parameters to the phage-attacked colonies

With the growth parameters determined, we use our model to fit the trajectories of our 20 phage-attacked colonies. Here we fit to match the visible bright-field radius of the colony to *V*_*t*_, and we match the region with a strong green fluorescence signal to *R*. Note that we here have no good measure of the uncertainties on colony radius for both the bright field and green fluorescence data and instead use the squared distance as our goodness-of-fit measure (See Materials and Methods for details).

The killing by phages in our model is described by two global parameters which both control the strength of the phage attack. These parameters are the phage penetration depth *γ* and the virulence parameter *ϵ*, which describes the phage killing rate in units of the bacterial growth rate. It should be noted that these two parameters somewhat compensate for each other, introducing the possibility of overfitting. Since the known latency time of phage T4 is comparable with the bacterial doubling time [28], we first show the result of fitting the value of *γ* with *ϵ* fixed to 1, and later discuss the fit when both are varied. We also allow the time of the phage invasion *T*_*i*_ to vary between each of the 20 data sets. Note that *T*_*i*_ represents the time at which the whole surface of the colony is penetrated down to the depth *γ*.

In Figure 5, we show the model fits to 5 of the 20 data sets with keeping *ϵ* = 1, which range from colonies that are quickly eliminated by the phages to colonies which seemingly retain an alive core of bacteria long after the phage attack begins. Model fits to all 20 data sets are included in Figure S4 in the supplement.

**Figure 5.**
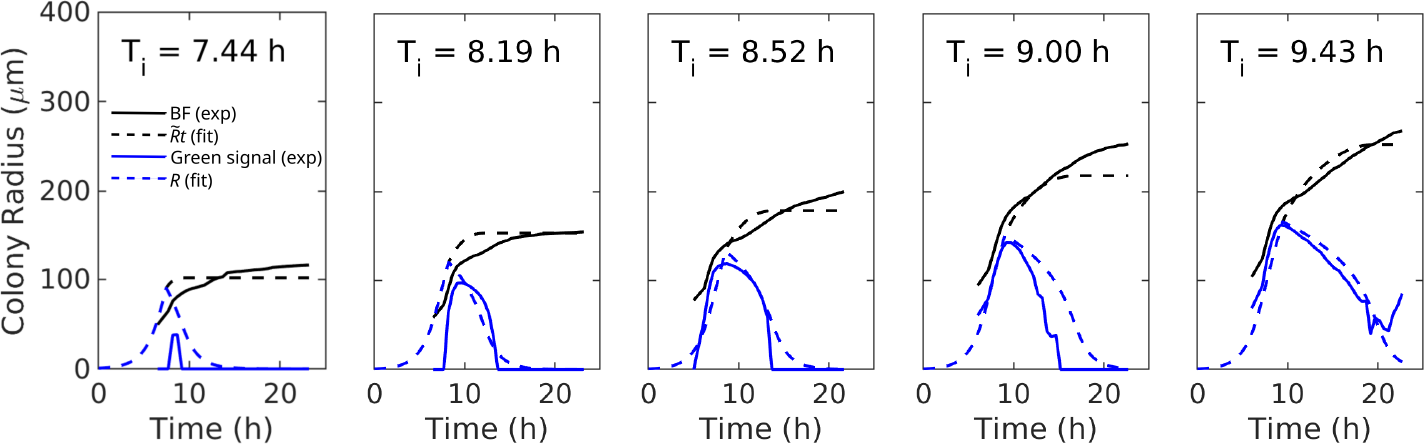
Killing by phages. We show 5 representative curves for the growth of *Escherichia coli* colonies in the presence of phage *T*4. Each panel shows a different colony attacked by phages at different times. The black lines indicate the time evolution of the visible radius of the colonies in bright-field conditions. The blue curves indicate the radius of the colony showing substantial green fluorescence, which we take as a proxy for alive biomass. The dashed lines indicate the least-squares fit of our model, with the predicted phage invasion time *T*_*i*_ shown on each graph. The blue dashed line shows the radius of the alive colony *R*, and the black dashed line shows the total radius including the dead biomass 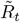. For each panel, the model uses the same values for the penetration depth*γ*= 27μm, and the phage virulence parameter is fixed to*ϵ*= 1, but different phage invation time *T*_*i*_. The results for all 20 colonies and fits are presented in Figure S4.

The fit gave the penetration depth *γ* = 27 μm. Assuming one cell layer is about 1 μm, this means 20-30 cell layers are penetrated (uncertainty is large due to elongated shape of *E. coli*). The fitted curve overestimated the decrease of the alive area at the start of the phage attack especially for small colonies/early phage attacks (Figure 5). This is likely because we assume that the phage starts to kill bacteria within the depth *γ* as soon as the phage attack initiates. In reality, it is more plausible that it takes a few phage generations for phage to penetrate many cell layers. When we allow both *γ* and *ϵ* to vary in the fitting, we have obtained *γ* = 3.5 μm and the phage virulence parameter *ϵ* = 9.4 as a fit. This relatively small *γ* compensated by an unrealistically large value of *ϵ* reproduced the fine structure in the time dependence of the killing curve better (Figure S5), suggesting the necessity of fine dynamics for reproducing the data better. The overall conclusion is that the *T*4 phage is able to penetrate quite deep into the colony eventually to efficiently kill bacteria.

We also tested if the large penetration depth is a robust conclusion against the choice of the threshold for green fluorescence signal to determine the alive radius. When we chose the threshold to be larger (smaller), we need to fit the dead cell and alive cell volume ratio *v* to be larger (smaller) than 1 (See materials and Method). The extracted alive radius and the fit allowing to change *v* from 1 with keeping *ϵ* = 1 are shown in the supplementary Fig. S6 for about 10% larger threshold and Fig. S7 for about 10% smaller threshold. The fitted value of *v* are about 1.3 and 0.7 and *γ* are about 30 μm and 25 μm, respectively. For the 10% smaller threshold, we start to see several colonies judged to be all dead once and then suddenly judged alive again because the fluctuating fluorescent signal crosses the threshold more than once. Therefore, the currently used or larger threshold will likely be more appropriate. Hence, the T4 penetration depth as deep as about 30 μm remains a robust conclusion of the current analysis.

The simple estimate of the critical colony radius *R*_*c*_ for colony survival (Equation (1)), which assumes that the growth rate does not change over time, predicts *R*_*c*_ ≈ 168 μm with the fitted parameter (*γ* = 27 μm with *ϵ* = 1). This colony size is reached only after about 8 to 9 hours of growth (Figure 4). However, the rate of colony expansion for an undisturbed bacterial colony significantly slows down around this time (Figure 6 blue line). To have a overview on expected growth and death of the colony, we simulated the alive radius over time for varying phage attack timing using the estimated parameters of *γ* = 27 μm with keeping *ϵ* = 1. The result is shown in Fig. 6. Here, we see that, if the infection happens about ∼10 hours or later, significant cells stay alive after 24 hours. Thus our model-based analysis of the data leaves a marginal possibility for long time survival of large colonies, as indeed also seen in the rightmost panel of Fig. 5.

**Figure 6.**
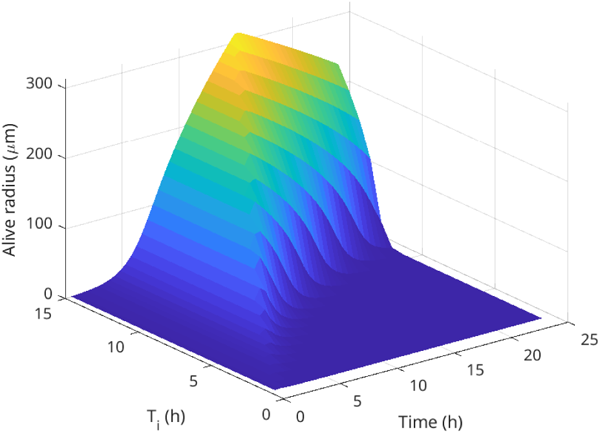
Expected response of the colony attacked at different timing. The simulated time evolution of the alive colony radius *R* using the estimated parameter for killing by phage (*γ*= 27μm with*ϵ*= 1) with systematically vaying the first phage attack timing.

## Discussion and Conclusion

Using an automated microscope, we tracked the dynamics of *Eschericia coli* microcolonies under attack by the virulent phage *T*4. This setup allowed us to monitor the fate of 20 different colonies growing in large wells and sprayed with phage after 5 to 7 hours of growth, corresponding to the fitted full-scale phage attack *T*_*i*_ at about 7 to 10 hours. By using an *Eschericia coli* variant that constitutively expresses green fluorescence protein, we were able to further estimate how the bacteria were being killed by the phages over time.

We developed a dynamical model to simulate the size of the bacterial colony when accounting for phages invading the colony’s surface. This model could fit the total and alive biomass to the experimental measurements. The model can generally mimic the trajectories of growth from the experiment. This allows us to extract critical parameters of the phage-bacteria dynamics. Most interestingly, we concluded that given the phage latency time is comparable with the bacterial generation time (*ϵ*= 1), the penetration depth of the phage *T*4 is as large as *γ*= 27μm.

The prediction of Ref. [11] was that bacterial colonies will, if they are sufficiently large, be able to outgrow the killing by phage, provided that the constant exponential growth can continue for surviving cells. For the virulent version of *P*1 phage used in Ref. [11], the experimentally observed critical size for colony outgrowth *R*_*c*_ about 25μm. Combined with eq. 1 and the *P*1 latency time [28] being about 60 min for *E. coli* growing with a generation time about 30 min, we estimate the penetration depth of phage *P*1*vir* was about 3μm. Compared to this, current model-based analysis suggests that, in addition to the latency time difference, the phage *T*4 is able to penetrate about 10-fold deeper into the colony. Therefore, the expected critical size for the colony to outgrow the killing by *T*4 becomes so large that the colony growth rate at that size has slowed down substantially due to limited nutrient supply. Since the host growth-state dependence of the phage growth parameter when the host is close to the stationary phase is still unknown [34, 36], our model cannot reliably predict the size of colonies that eventually may survive a phage attack.

We do not have an explanation for why phage *T4* can invade the colony so effectively. One simple possibility is the colony is less dense and has more pores for phage to diffuse than we expect. However, the bacteria colonies were grown in the same concentration of soft agar in the experiments with *T4* and those *P*1_*vir*_ [11]. Hence, it is more likely that it is a phage-dependent property. It is expected that deeper penetration is easier if the phages adsorb less to bacteria [11], but the adsorption rates are not significantly different between the two phages [28]. Further experimental studies to visualize phage penetration dynamics inside a colony are desirable, for example, using laser sheet microscopy.

Before concluding, we here discuss a few factors that were not taken into account in the current model.

First, the model does not consider the appearance of phage-resistant mutants, which happens typically by the loss of phage receptors. For the phage T4, the reported mutation rate for resistance is 8 to 50 × 10 ^*−*8^ per bacterial generation time [41]. This means that the resistant cells will be frequently observed only after the colony has reached 10^8^ cells, i.e., 300μm in radius. Hence, it will become a significant effect if the phage attack happens after 12h growth or later (see Fig. 4). Our current analysis was focused on phage attacks on smaller colonies, where resistance should occur rarely. In addition, if the rare event of early mutation happens and the resistant cell number increases significantly before the phage attack, they will be visible because their progenies will be in physical proximity and stay fluorescent after the phage attack, as shown in Fig. 3. In the current analysis, we excluded those colonies. It would be an exciting extension to include the resistant bacteria. However, such a model requires stochastic and a fully three-dimensional description since the appearance of the resistant cell locally breaks the spherical symmetry of a colony.

Second, *T*4 phage is known to show lysis inhibition, where the superinfection of an already T4-infected cell causes a delay in lysis [42, 43, 44]. Our model does not include burst events explicitly but assumes that an infected cell lyses after a certain time scale, and that determines the advancement of the infected layer of thickness *γ*. While it is likely that there is a high superinfection on the surface to the middle of the infected layer, the number of phages that reach deeper should decay exponentially and at the front of the infection, i.e., at the deepest point in the colony where a phage has reached, it is likely that the infection happens at multiplicity one. Hence, it is likely that the burst time scale to invade the colony is not affected by lysis inhibition. It is interesting to note that, even though the clear region of T4 plaques is quite small, it has been proposed that the infection front of a T4 plaque actually extends as far as those of lysis inhibition mutants that show larger clear region [44].

In conclusion, we analyzed colony-level protection from attack by the virulent phage *T4*, and found that this phage was surprisingly efficient in dissolving even large colonies by penetrating many cell layers. This striking ability of *T*4 differs from the much smaller penetration of phage *P*1_*vir*_, and thereby suggests a new type of phage-specific infection strategy under high bacterial densities. The deep penetration of *T*4 substantially reduces colony-level protection. However, a large colony provides some shielding and substantially delays the lysis in the centre of the colony making it possible that some very large colonies may show long time survival of T4 attack.

## Author Contributions

RSE, FL, SLS, KS, and NM designed and performed research. RSE developed the software. RSE and NM analyzed data. RSE, SLS, KS, and NM wrote the manuscript.

## Acknowledgement

SLS was supported by Independent Research Fund Denmark (grant 8049-00071B). NM was supported by Novo Nordisk Foundation (NNF21OC0068775).

