## Supplementary material for "The dynamics of phage predation on a microcolony"

### Supplementary Material for:
### The dynamics of phage predation on a microcolony

Rasmus Skytte Eriksen[1,2], Frej Larsen[3,4],
Sine Lo Svenningsen[3], Kim Sneppen[1], and Namiko Mitarai[1]

[1] The Niels Bohr Institute, University of Copenhagen, Copenhagen, Denmark

[2] Department of Epidemiology Research, Statens Serum Institut, Copenhagen, Denmark

[3] Department of Biology, University of Copenhagen, Copenhagen, Denmark.

[4] Department of food science, University of Copenhagen, Copenhagen, Denmark.

October 23, 2023

#### Contents

**S1 Laplacian in spherical coordinates on a non-equidistant grid** **2**

**S2 Green fluorescence threshold value** **4**

**S3 Outliers** **5**

**S4 All model fits** **5**

**S5 Fit with varying both $\epsilon$ and $\gamma$** **7**

**S6 A higher GFP threshold to determine the alive radius** **8**

**S7 A lower GFP threshold to determine the alive radius** **9**

#### Parameter values used in simulations

| Name | Value | Description |
|---|---|---|
| $r_{\max}$ | 2000 µm | Size of the simulated space |
| $\Delta r$ | 0.05 µm | Length scale of the lattice |
| $k$ | 1.02 | Scaling factor for the lattice |
| $n_0$ | $(4.68 \pm 0.09) \cdot 10^{-2}$ | Initial density of nutrients (units of reproduction) |
| $D$ | $(1.29 \pm 0.03) \cdot 10^{5}$ µm$^2$/h | Diffusion constant for the nutrient |
| $K$ | $(2.75 \pm 0.06) \cdot 10^{-2}$ | Michaelis-Menten half-speed constant |
| $\mu g_{\max}$ | $(3.33 \pm 0.012)$ h$^{-1}$ | Maximal growth rate |
| $\gamma$ | 27 µm | Phage penetration depth (fitted with killing by phage multiplier $\epsilon$ fixed to 1) |

Table S1: Parameter values used / fitted in the model.

#### S1 Laplacian in spherical coordinates on a non-equidistant grid

Since the distance between the nodes in the grid is non-uniform we use Neville's algorithm to define the second order polynomial which intersects the values of three consecutive nodes.

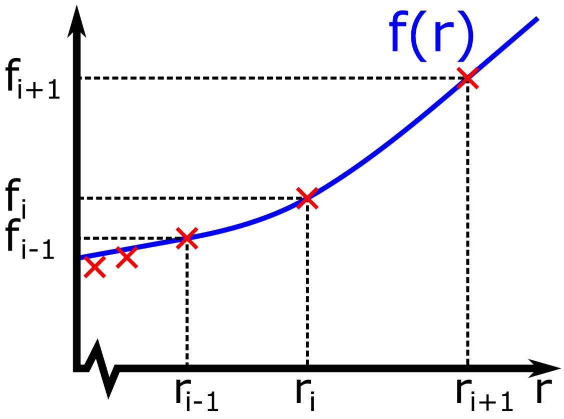


Figure S1: Discretization scheme.

The polynomial has the following form:

$$
\begin{aligned}
f(r) &= \frac{(r-r_i)(r-r_{i+1})f_{i-1}}{(r_{i-1}-r_i)(r_{i-1}-r_{i+1})} \\
&+ \frac{(r-r_{i-1})(r-r_{i+1})f_i}{(r_i-r_{i-1})(r_i-r_{i+1})} \\
&+ \frac{(r-r_i)(r-r_{i-1})f_{i+1}}{(r_{i+1}-r_i)(r_{i+1}-r_{i-1})}
\end{aligned}
$$

Since the problem has spherical symmetry, the laplacian $\nabla^2$ takes a simplified form in spherical coordinates:

$$\nabla^2 = \frac{1}{r^2}\frac{d}{dr}\left(r^2\frac{d}{dr}\right)$$

We can combine the laplacian with the result derived from Neville's algorithm to approximate the laplacian on the internal nodes:

$$
\begin{aligned}
\nabla^2 f(r_i) &= \frac{1}{r^2}\frac{d}{dr}\left(r^2\frac{d}{dr}\right)f(r)\Big|_{r=r_i} \\
&= \frac{1}{r^2}\frac{d}{dr}r^2\left(\frac{(2r-r_i-r_{i+1})f_{i-1}}{(r_{i-1}-r_i)(r_{i-1}-r_{i+1})}+\frac{(2r-r_{i-1}-r_{i+1})f_i}{(r_i-r_{i-1})(r_i-r_{i+1})}+\frac{(2r-r_i-r_{i-1})f_{i+1}}{(r_{i+1}-r_i)(r_{i+1}-r_{i-1})}\right)\Bigg|_{r=r_i} \\
&= \frac{2}{r}\left(\frac{(3r-r_i-r_{i+1})f_{i-1}}{(r_{i-1}-r_i)(r_{i-1}-r_{i+1})}+\frac{(3r-r_{i-1}-r_{i+1})f_i}{(r_i-r_{i-1})(r_i-r_{i+1})}+\frac{(3r-r_i-r_{i-1})f_{i+1}}{(r_{i+1}-r_i)(r_{i+1}-r_{i-1})}\right)\Bigg|_{r=r_i}
\end{aligned}
$$

$$\nabla^2 f(r_i) = \frac{2}{r_i}\left(\frac{(2r_i-r_{i+1})f_{i-1}}{(r_{i-1}-r_i)(r_{i-1}-r_{i+1})}+\frac{(3r_i-r_{i-1}-r_{i+1})f_i}{(r_i-r_{i-1})(r_i-r_{i+1})}+\frac{(2r_i-r_{i-1})f_{i+1}}{(r_{i+1}-r_i)(r_{i+1}-r_{i-1})}\right) \tag{S1}$$

Special care must be taken at the nodes $r_0$ and $r_N$. For the node at $r=0$, we take the limit of the laplacian:

$$\nabla^2 f(0) = \lim_{r\to 0}\left(\frac{1}{r^2}\frac{d}{dr}\left(r^2\frac{d}{dr}\right)f(r)\right)$$

We use the identity: $\frac{d}{dr}(r^2\frac{d}{dr}) = 2r\frac{d}{dr} + r^2\frac{d^2}{dr^2}$

$$
\begin{aligned}
\nabla^2 f(0) &= \lim_{r\to 0}\left(\frac{1}{r^2}\left(2r\frac{d}{dr}+r^2\frac{d^2}{dr^2}\right)f(r)\right)\\
&= \lim_{r\to 0}\left(\frac{2}{r}\frac{d}{dr}f(r)\right)+\frac{d^2}{dr^2}f(0)\\
&= 2\frac{d^2}{dr^2}f(0)+\frac{d^2}{dr^2}f(0)\\
&= 3\frac{d^2}{dr^2}f(0)
\end{aligned}
$$

Again we input the result from Neville's algorithm to get our numerical approximation:

$$
\begin{aligned}
\nabla^2 f(0) &= 3\frac{d^2}{dr^2}f(0)\\
&= \frac{6f_{-1}}{(r_{-1}-r_0)(r_{-1}-r_1)}+\frac{6f_0}{(r_0-r_{-1})(r_0-r_1)}+\frac{6f_1}{(r_1-r_0)(r_1-r_{-1})}
\end{aligned}
$$

Here we invoke the Neumann boundary condition by treating the node $r_{-1}$ as a ghost node with value $f_{-1} = f_1$ and located at $r_{-1} = -r_1$

$$
\begin{aligned}
\nabla^2 f(0) &= \frac{6f_1}{(-r_1)(-r_1-r_1)}+\frac{6f_0}{r_1(-r_1)}+\frac{6f_1}{r_1(r_1+r_1)}\\
&= \frac{6f_1}{2r_1^2}-\frac{6f_0}{r_1^2}+\frac{6f_1}{2r_1^2}
\end{aligned}
$$

$$
\nabla^2 f(0) = \frac{6}{r_1^2}\left(f_1-f_0\right) \tag{S2}
$$

Finally we need to consider the end node $r_N$: As with the node at $r = 0$, we use the Nuemann boundary condition and include a ghost node at $r_{N+1} = r_N + (r_N - r_{N-1})$ with the value $f_{N+1} = f_{N-1}$. Inserting the ghost node explicitly into (S1) yields:

$$
\begin{aligned}
\nabla^2 f(r_N) &= \frac{2}{r_N}\left(\frac{r_{N-1}f_{N-1}}{2(r_{N-1}-r_N)r_N}+\frac{(2r_{N-1}-r_N)f_N}{(r_N-r_{N-1})(r_N+r_{N-1})}+\frac{(2r_N-r_{N-1})f_{N-1}}{2(r_N-r_{N-1})(r_N-r_{N-1})}\right)\\
&= \frac{2}{r_N}\left(\frac{r_{N-1}f_{N-1}}{2(r_{N-1}-r_N)^2}-\frac{r_Nf_N}{(r_N-r_{N-1})^2}+\frac{(2r_N-r_{N-1})f_{N-1}}{2(r_N-r_{N-1})^2}\right)
\end{aligned}
$$

$$
\nabla^2 f(r_N) = \frac{2}{(r_N-r_{N-1})^2}(f_{N-1}-f_N) \tag{S3}
$$

#### S2 Green fluorescence threshold value

The principle behind our image analysis is to use a simple threshold to determine the rough outline of the bacterial colonies. For the bright-field images, this threshold value can be chosen over a wide interval since the contrast between the colony and the medium is large even at the colony edge. However, the image analysis of our images of the green fluorescence protein (GFP) activity in the bacterial colony is on the other hand very sensitive to the chosen threshold value. This GFP signal essentially forms a gradient radiating out from the centre of the colony and tapering off towards the edges of the colony. We here use our model to estimate the appropriate threshold value for the algorithm. We do so by sweeping through a range of threshold values and comparing the density of the alive bacteria with the dead bacteria (see main text for details). This density is expressed via the transparency parameter $\nu$ which we argue should take a value of $\sim 1$. In Figure S2, we plot the estimated value of $\nu$ as a function of the threshold value and find that a threshold value of 1.8 satisfies the requirement $\nu \sim 1$.

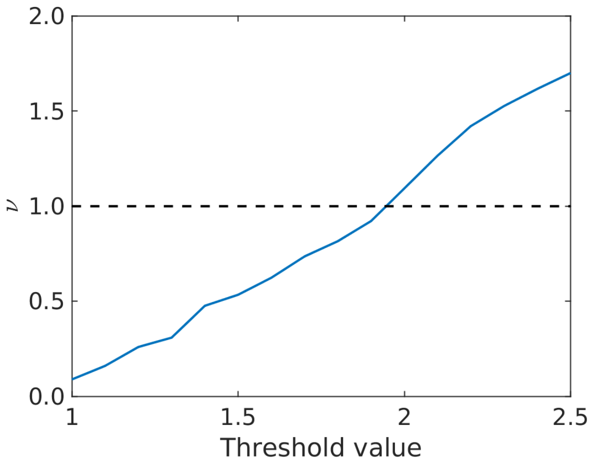


Figure S2: **Image analysis threshold.** We plot the transparency parameter $\nu$ as a function of the threshold value used in the image analysis. This parameter reflects the relative density of alive bacteria and dead bacteria.

#### S3 Outliers

After running our image analysis algorithm, we obtain 5 growth trajectories which we classify as outliers. Here, 3 of these trajectories seemingly correspond to the emergence of resistant mutants where the green fluorescence signal initially drops when the phage attack begins, but, after enough time, this signal increases again. One trajectory does not show this tell-tale sign of resistance, but rather show a seemingly bimodal killing curve. The last trajectory is unique in that the green fluorescence signal cannot be detected.

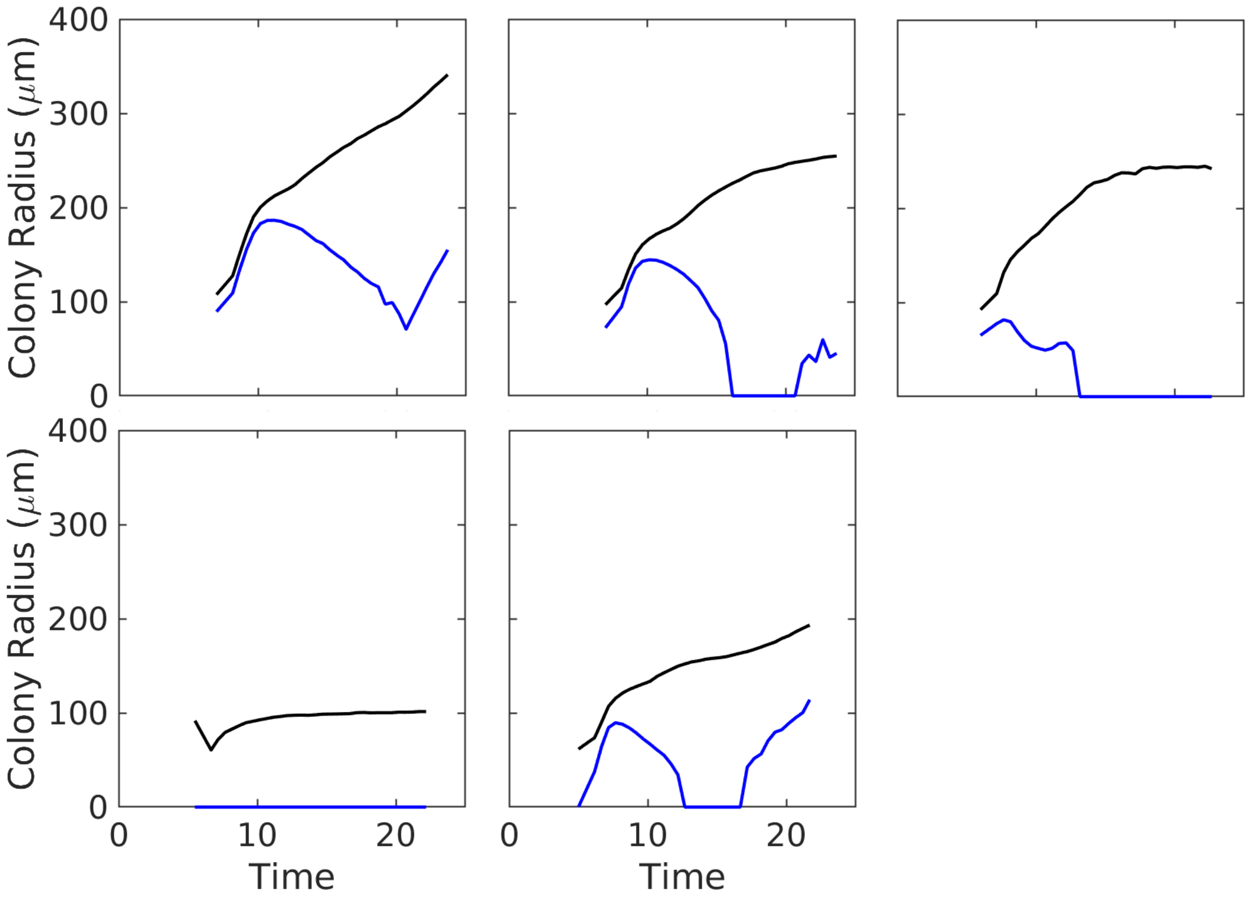


Figure S3: **Outlier trajectories.** Our analysis provides us with 5 growth trajectories which qualitatively differs from the majority data sets. These include three trajectories which show a secondary growth which is consistent with the emergence of resistant mutants. One trajectory has an anomalous bi-modal growth curve. The green fluorescence signal from one colony is not registered.

#### S4 All model fits

In total, we have 20 trajectories for colony growth that we include in our analysis. We fit our model to this entire set of 20 trajectories simultaneously. This is achieved by allowing the phage invasion time to vary independently for each data set but with the value for the phage penetration depth $\gamma$ fixed across all data sets during the fit. We show our model fit with $\epsilon = 1$ (fixed during the fit) $\gamma = 27\mu$m (obtained from fit) to all 20 data sets in Figure S4. These are sorted by the estimated phage invasion time $T_i$ ranging from smallest to largest.

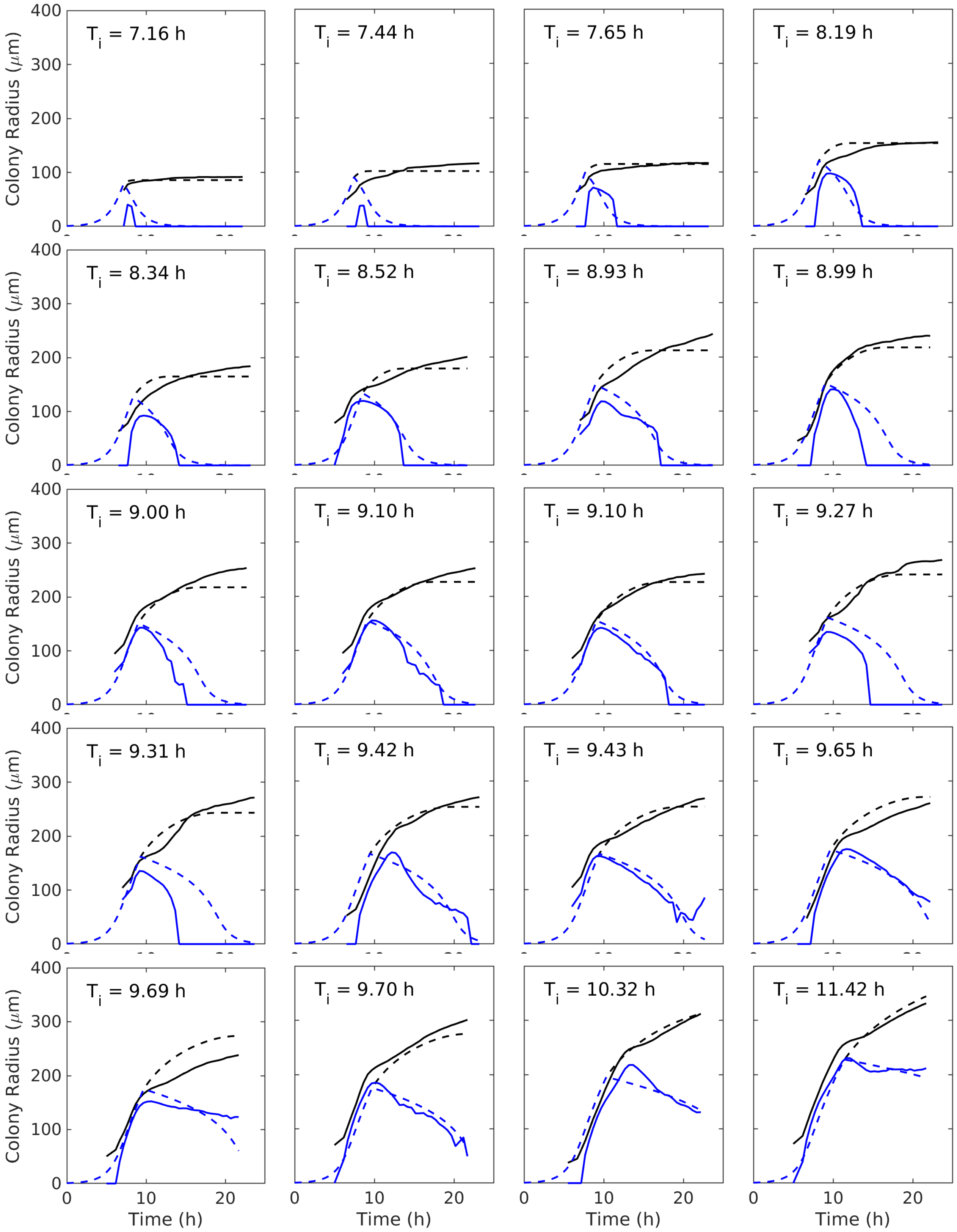


Figure S4: **All model fits with** $\epsilon = 1$**.** We show the model-fits to the 20 different colony trajectories extracted from our experiment, with $\epsilon = 1$. This results in the fitted value of $\gamma = 27\mu$m. Bright-field colony radius is shown as solid black lines, with the model prediction shown as the corresponding dashed line. The signal from the green fluorescence protein is shown as the solid blue line, and the model prediction for the radius of the alive bacteria is shown as the dashed blue line. Some of the panels are replications of those in Fig. 5.

#### S5 Fit with varying both $\epsilon$ and $\gamma$

Figure S5 shows the fit obtained by allowing both $\epsilon$ and $\gamma$ to vary, which gives a better fit but an unrealistically large value of $\epsilon = 9.4$ with small $\gamma = 3.5$ μm.

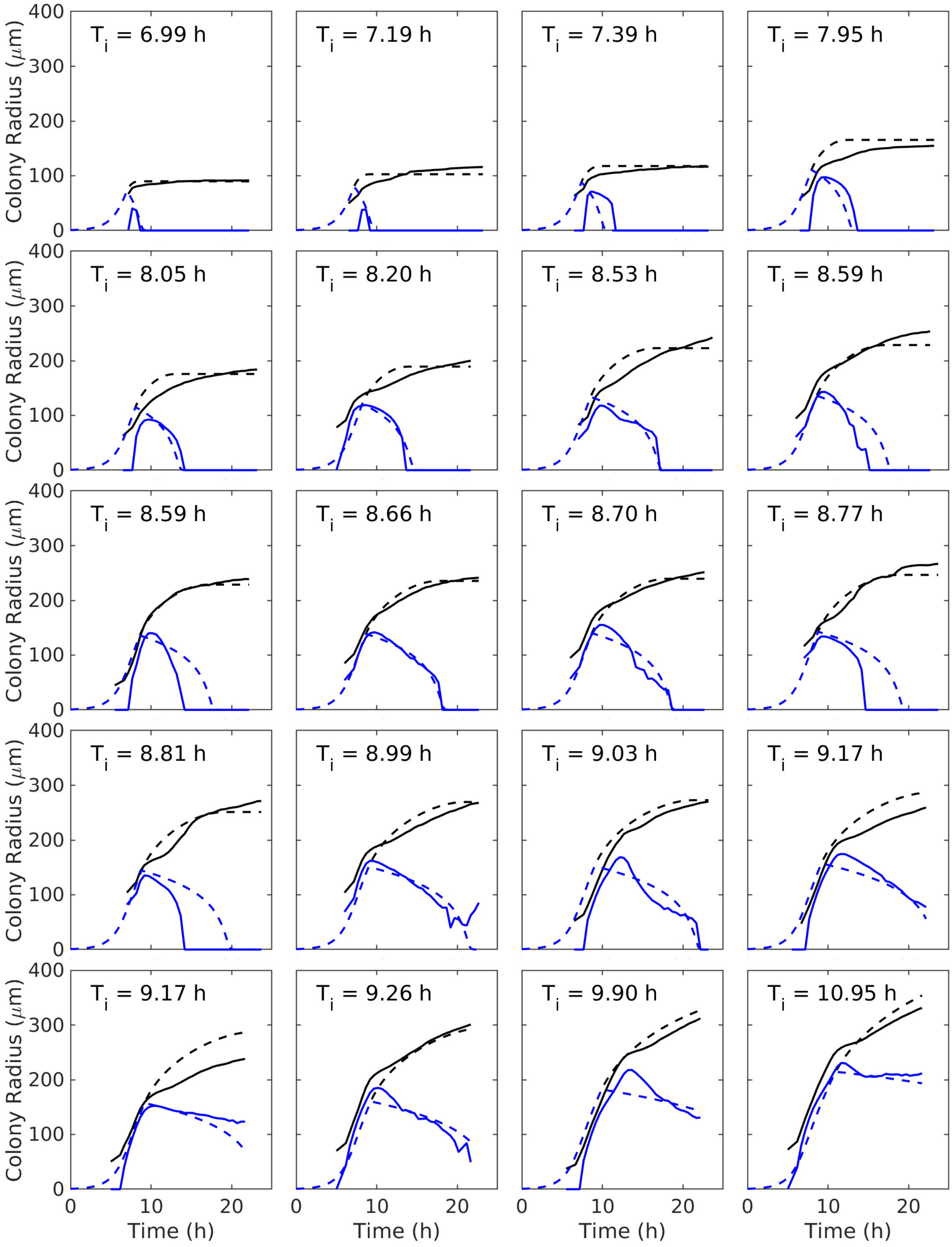


Figure S5: **All model fits.** Simlar to Fig. S4, we show the model-fits to the 20 different colony trajectories extracted from our experiment with $\epsilon = 9.4$ and $\gamma = 3.5$ μm. The bright-field colony radius is shown as solid black lines with the model prediction shown as the corresponding dashed line. The signal from the green fluorescence protein is shown as the solid blue line and the model prediction for the radius of the alive bacteria shown as the dashed blue line.

#### S6 A higher GFP threshold to determine the alive radius

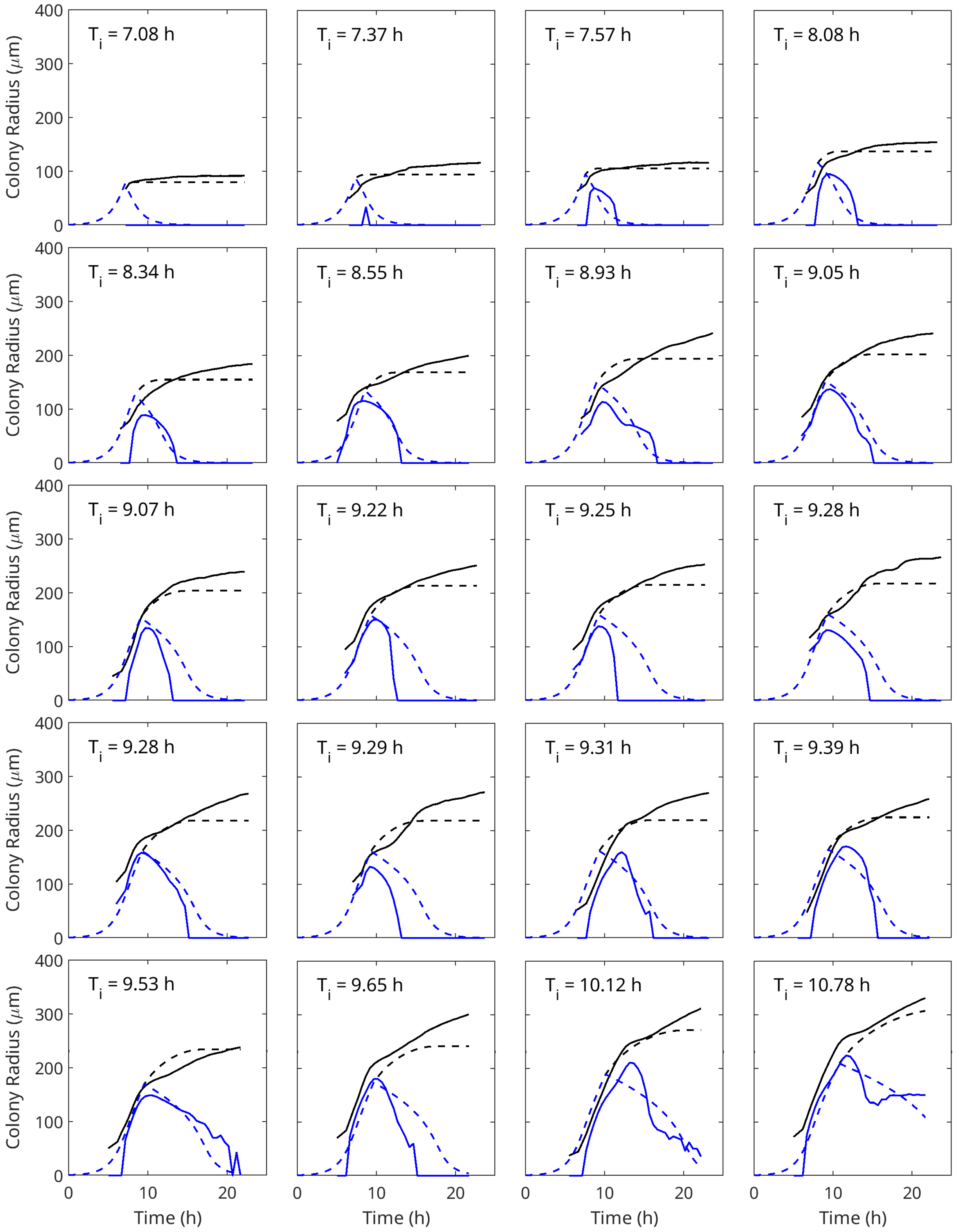


Figure S6: **Data and model fits with the threshold value for image analysis to be 2.0.** In the fit, $\epsilon$ was set to 1, and the transparency parameter $\nu$ and the penetration radius $\gamma$ was fitted to the data. We obtained $\nu = 1.3$ and $\gamma = 30$ μm as the best fit.

#### S7 A lower GFP threshold to determine the alive radius

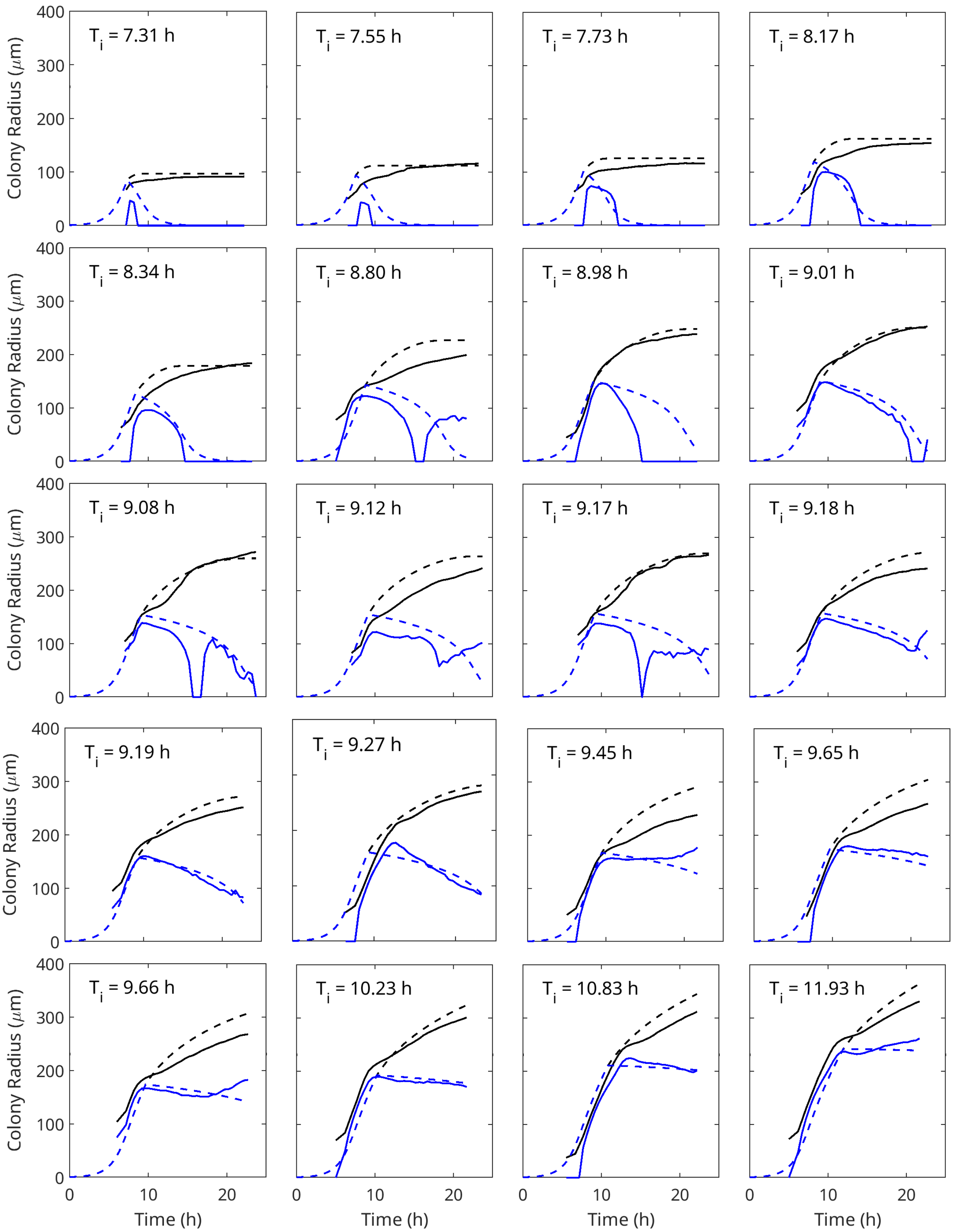


Figure S7: **Data and model fits with the threshold value for image analysis to be 1.6.** In the fit, $\epsilon$ was set to 1, and the transparency parameter $\nu$ and the penetration radius $\gamma$ was fitted to the data. We obtained $\nu = 0.72$ and $\gamma = 25$ μm as the best fit.
